# Four new *Scopalina* from Southern California: the first Scopalinida (Porifera: Demospongiae) from the temperate Eastern Pacific

**DOI:** 10.1101/2020.08.11.246710

**Authors:** Thomas L. Turner

## Abstract

Sponges (phylum Porifera) are common inhabitants of kelp forest ecosystems in California, but their diversity and ecological importance are poorly characterized in this biome. Here I use freshly collected samples to describe the diversity of the order Scopalinida in California. Though previously unknown in the region, four new species are described here: *Scopalina nausicae* **sp. nov**., *S. kuyamu* **sp. nov**., *S. goletensis* **sp. nov**., and *S. jali* **sp. nov.**. These discoveries illustrate the considerable uncharacterized sponge diversity remaining in California kelp forests, and the utility of SCUBA-based collection to improve our understanding of this diversity.

## Introduction

The order Scopalinida is young: it was created in 2015 (Morrow & Cárdenas 2015). The independent evolution of this lineage, however, is old. Phylogenies based on ribosomal DNA place the order, together with the freshwater sponges, as the sister clade to all other extant orders in the subclass Heteroscleromopha (Morrow *et al*. 2013). This interesting phylogenetic position motivates further investigation of this group. The order is comprised of only two genera, *Scopalina* and *Svenzea*, though genetic data suggest *Stylissa flabelliformis* may also be in the order (other genotyped *Stylissa* cluster with the order Agelisida (Erpenbeck *et al*. 2006; Morrow *et al*. 2012; Morrow & Cárdenas 2015). Scopalinida have been found to be sources for novel anti-microbial or anti-tumor compounds, which further motivates efforts to characterize their diversity (Avilés *et al*. 2013; Vicente *et al*. 2015; Wei *et al*. 2007).

Nearly all Scopalinida are described from warm waters in the Mediterranean, Caribbean, South-West Pacific, and Madagascar (van Soest *et al*. 2019). The only previous exceptions to this pattern are the two most recently described species, from the Falkland Islands, which were found by hand-collecting sponges while SCUBA diving (Goodwin *et al*. 2011). Collecting by hand has been shown to be a productive way to discover new sponges from rocky areas in the shallow subtidal (Goodwin *et al*. 2011; Goodwin & Picton 2009), but past sponge surveys in Southern California have primarily been conducted via dredging or by collecting in the intertidal zone (Bakus & Green 1987; Green & Bakus 1994; de Laubenfels 1932; Sim & Bakus 1986). As a result, some common sponges found in California kelp forests — which occur on shallow hard-bottom substrate — remain unknown to science. Kelp forests in California are experiencing rapid changes due to anthropogenic impacts, and considerable work is focused on understanding kelp forest ecology to better predict and/or mitigate these impacts (Caselle *et al*. 2018; Castorini *et al*. 2018; Eger *et al*. 2020; Miller *et al*. 2018; Reed *et al*. 2016). The roles that sponges play in this ecosystem are unknown, but describing the species composition of the system represents a first and necessary step to improving this understanding. To better understand the abundance and distribution of shallow-water marine sponges in California, I have used SCUBA to collect over 300 individuals, mostly from kelp forest habitats in the Santa Barbara Channel. I have previously used this collection to revise the order Tethyida in California (Turner 2020b), but the other sponges remain to be described. Ten of these new samples can be assigned to the order Scopalinida, and are described herein as four new species in the genus *Scopalina*.

## Methods

### Collections

I collected sponge samples by hand with a knife. Each sample was placed immediately in a plastic bag with copious seawater. After the dive, these bags were put on ice for 2-12 hours. Samples were then preserved in 95% ethanol, which was exchanged for new preservative after 1-3 days, and changed again if it remained cloudy. Samples were vouchered with the California Academy of Sciences in San Francisco; archival information is listed in table 1.

**Table 1.** Archival information for vouchered samples and samples with sequence data. CASIZ: California Academy of Sciences Invertebrate Zoology.

| Collection location | Lat | Long | Date | Species | Collection ID | Museum voucher | CO1 sequence | 28S sequence |
| --- | --- | --- | --- | --- | --- | --- | --- | --- |
| Big Rock, Santa Cruz Island, CA | 34.05220 | -119.57360 | 1/19/20 | *Scopalina jali* | TLT567 | CASIZ 235466 | - | MT586556 |
| Naples Reef, Santa Barbara, CA | 34.42212 | -119.95154 | 9/26/19 | *Scopalina jali* | TLT324 | CASIZ 235467 | MT583199 | MT586558 |
| Naples Reef, Santa Barbara, CA | 34.42212 | -119.95154 | 12/10/19 | *Scopalina jali* | TLT558 | CASIZ 235468 | - | - |
| Naples Reef, Santa Barbara, CA | 34.42212 | -119.95154 | 7/31/19 | *Scopalina kuyamu* | TLT122 | CASIZ 235469 | MT583200 | MT586559 |
| Elwood Reef, Santa Barbara, CA | 34.41775 | -119.90150 | 10/23/19 | *Scopalina goletensis* | TLT307 | CASIZ 235470 | MT583198 | MT586557 |
| Coal Oil Point, Santa Barbara, CA | 34.40450 | -119.87890 | 8/30/19 | *Scopalina nausicae* | TLT341 | CASIZ 235471 | - | MT586560 |
| Isla Vista Reef, Santa Barbara, CA | 34.40278 | -119.85755 | 8/1/19 | *Scopalina nausicae* | TLT132 | CASIZ 235472 | - | MT586561 |
| Arroyo Quemado Reef, Santa Barbara, CA | 34.46775 | -120.11905 | 1/7/20 | *Scopalina nausicae* | TLT549 | CASIZ 235473 | - | - |
| Goalpost, Point Loma, San Diego, CA | 32.69438 | -117.26860 | 2/7/20 | *Scopalina nausicae* | TLT471 | CASIZ 235474 | MW834351 | MT586562 |

Samples were photographed *in situ* with an Olympus TG5 before collection. I photographed all sponge morphotypes found at each site, so that presence/absence across sites could be used to form hypotheses about sponge distributions and habitat. Table 2 contains information about all locations investigated, together with an estimate of search effort at each location. These locations include intertidal sites, floating docks in marinas, and subtidal sites searched on SCUBA. Subtidal sites were generally shallow (<15 m) rocky reefs, but a few sites were deeper (up to 27 m) and include artificial reefs and oil rigs. Scopalinida were only found at shallow, subtidal, natural reefs. Field photos of all sponges have been archived with vouchers, and also posted as searchable, georeferenced records on the site iNaturalist.org.

**Table 2.**
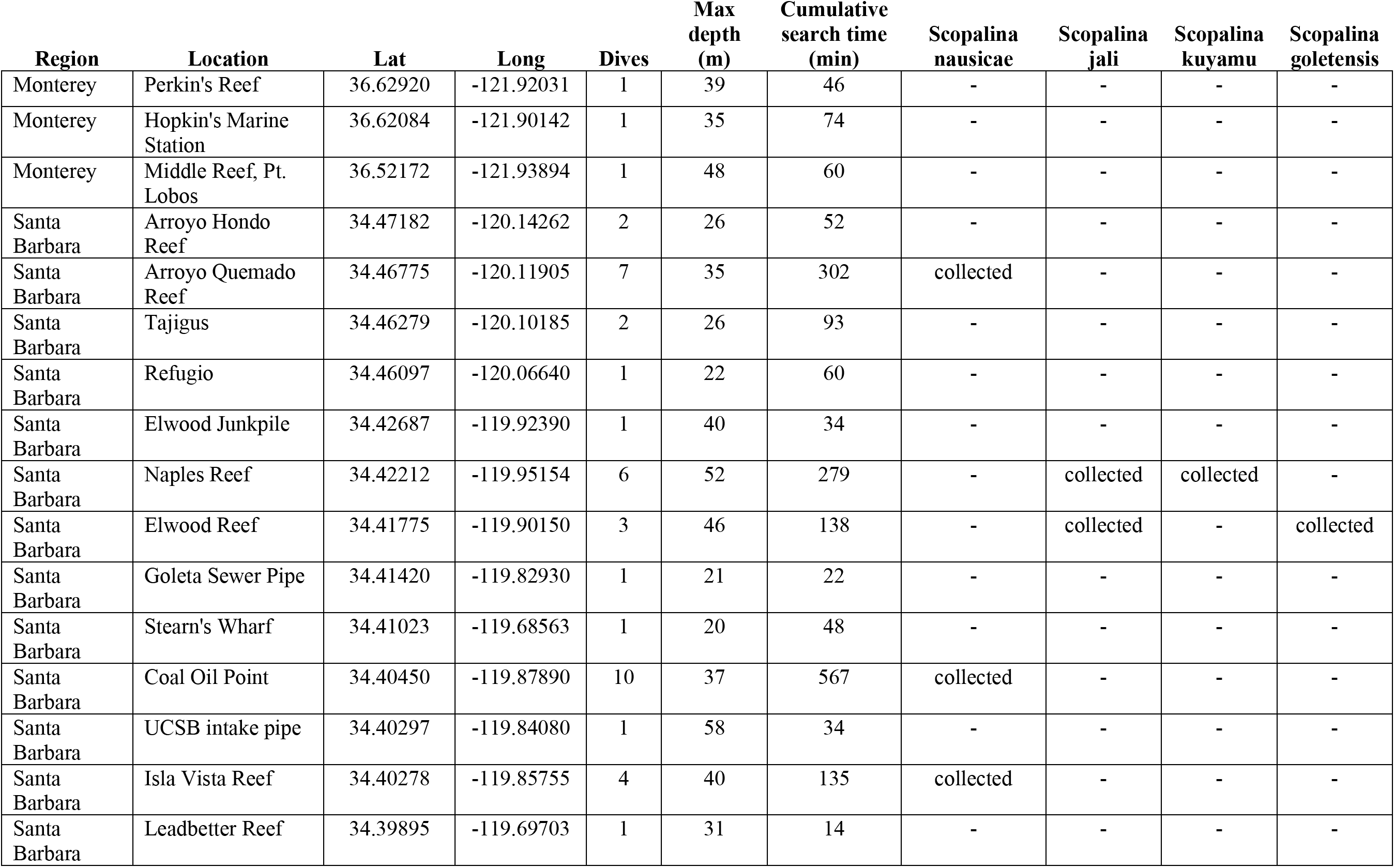
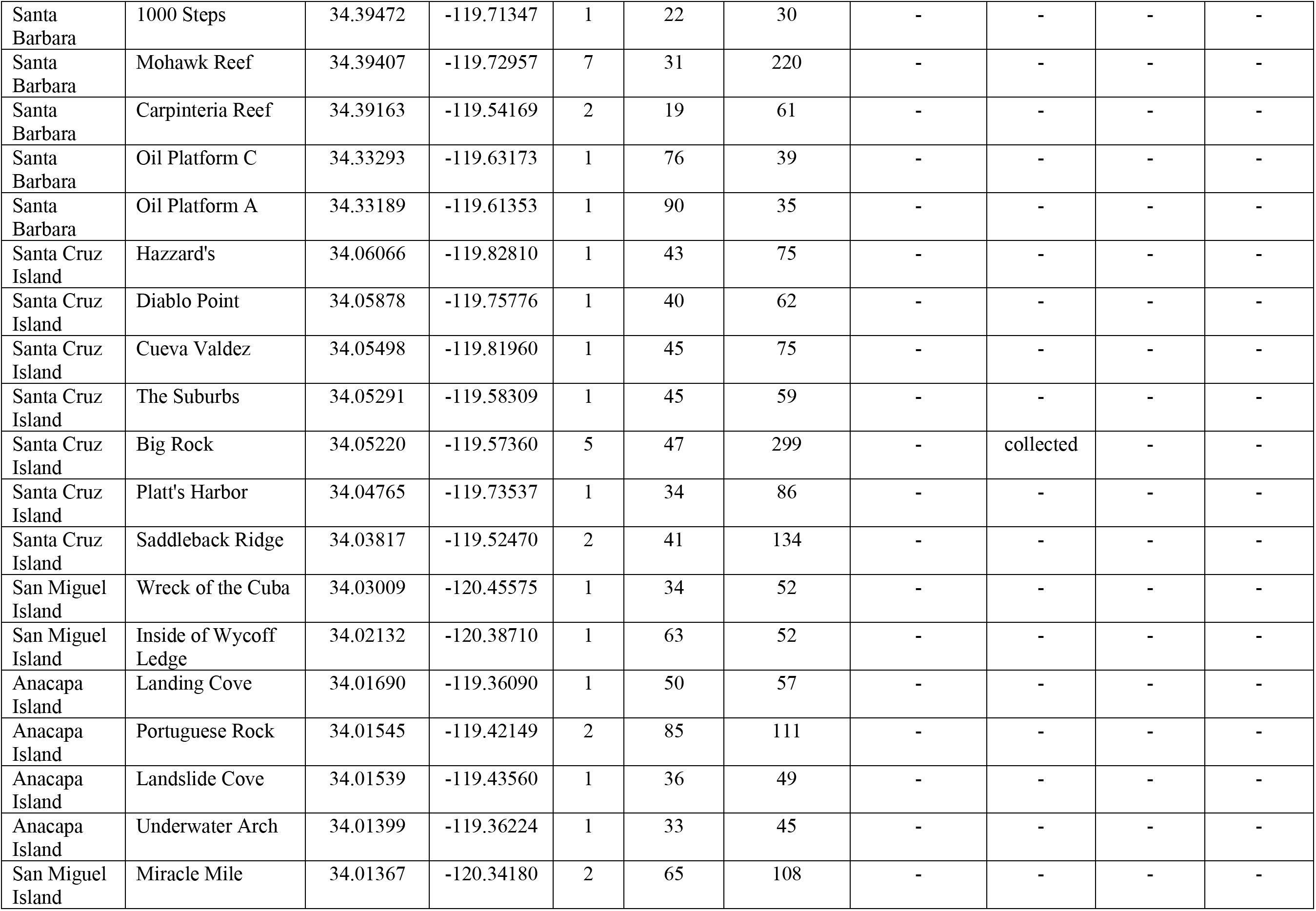
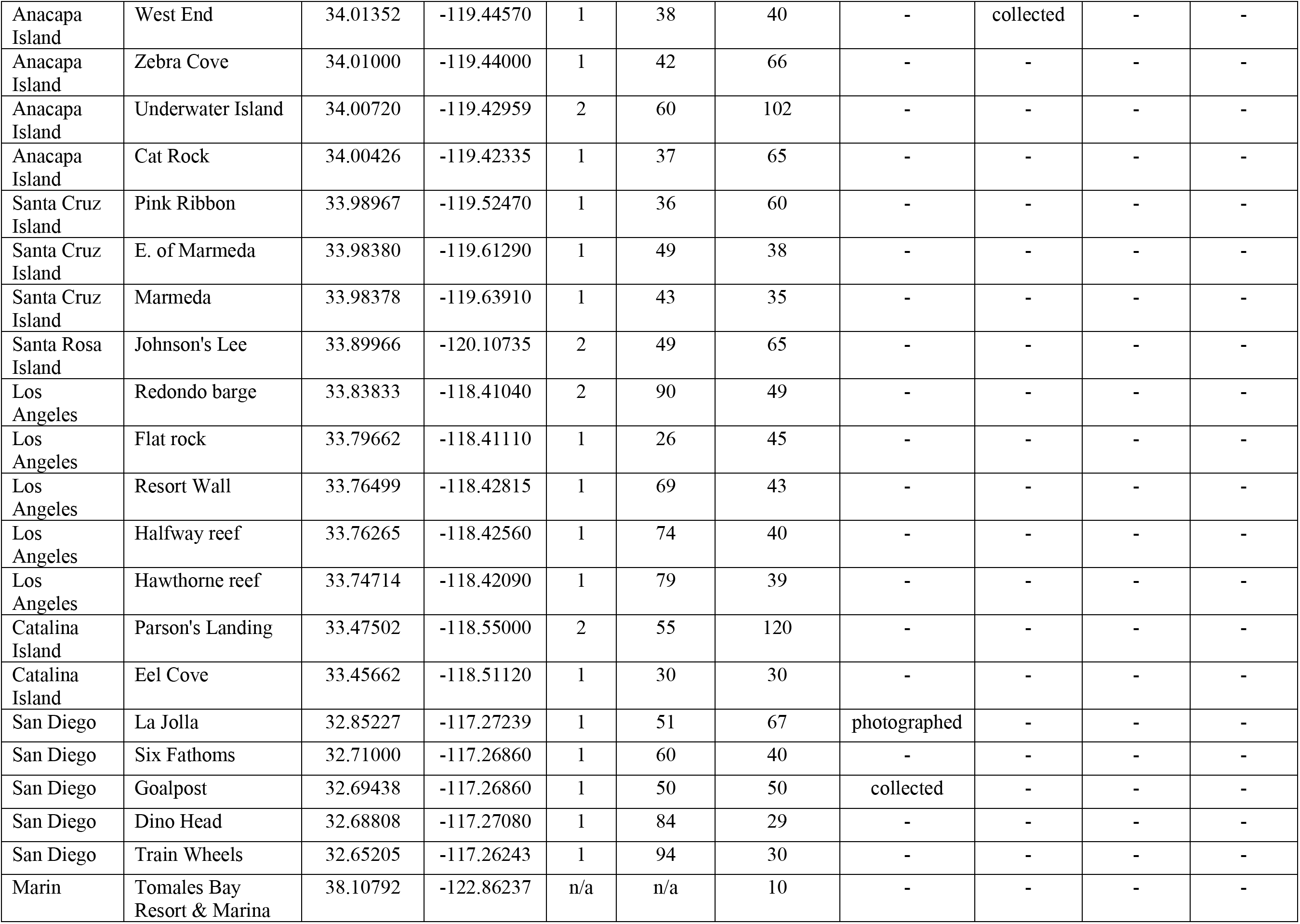
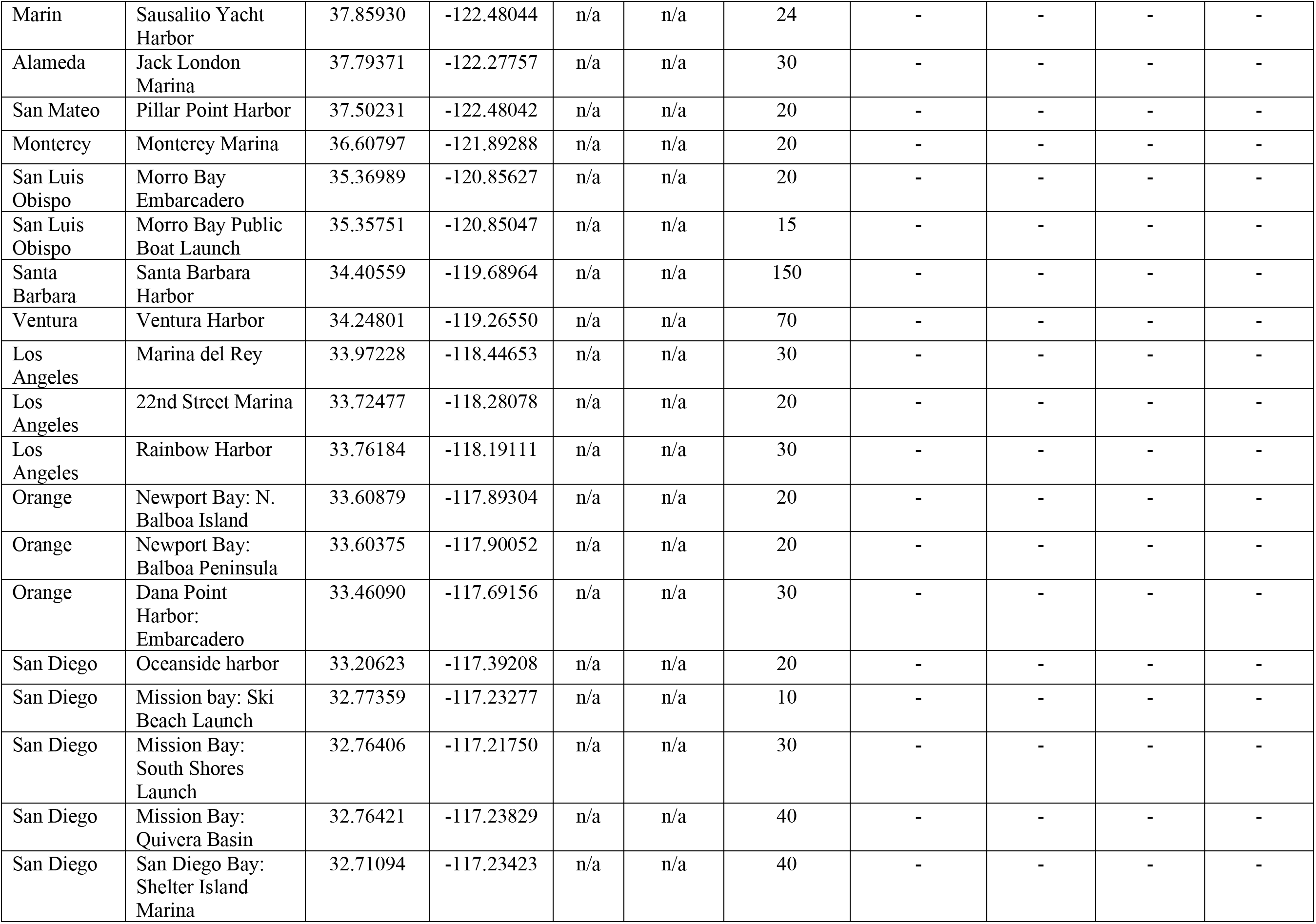
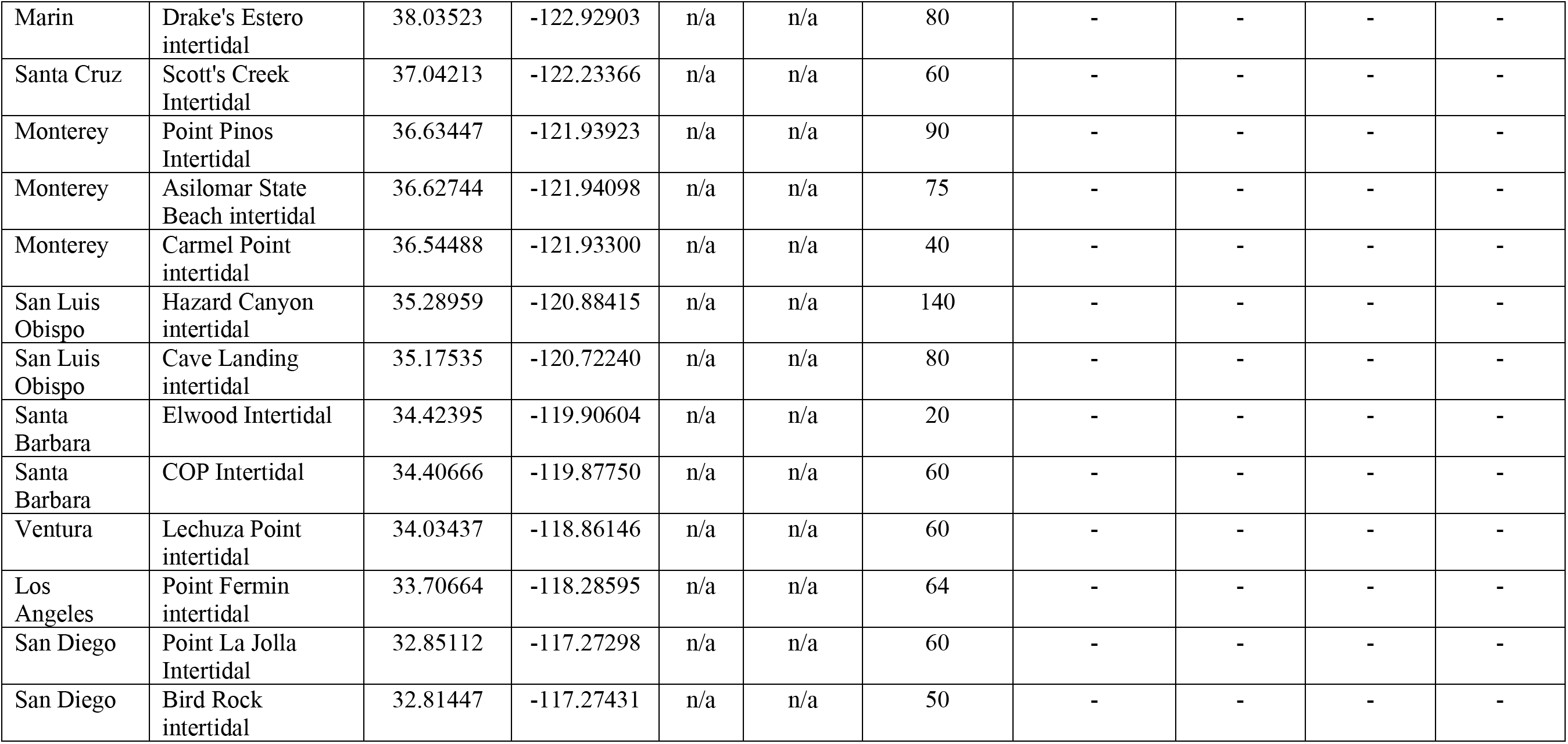
Sampling effort and locations. Harbor and intertidal sites are listed last. For subtidal sites, the number of sampling dives is listed, and the cumulative dive time as an estimate of search effort.

### Spicules

Spicule preparations were performed by digesting soft tissue subsamples in bleach. With the spicules settled at the bottom of the reaction tube, the bleach was then pipetted off and replaced with distilled water; this was repeated several times. Spicules were imaged using a D3500 SLR camera (Nikon) with a NDPL-1 microscope adaptor (Amscope) attached to a compound triocular microscope. A calibration slide was used to determine the number of pixels per mm, and spicules were then measured using ImageJ (Schneider *et al*. 2012). Spicule length was measured as the longest possible straight line from tip to tip, even when spicules were curved or bent. Spicule width was measured at the widest point. To image spicular architecture, I hand-cut perpendicular sections and, when possible, removed sections of ectosome. Sections were digested in a mixture of 97% Nuclei Lysis Solution (Promega; from the Wizard DNA isolation kit) and 3% 20mg/ml Proteinase K (Promega). This digestion eliminates cellular material while leaving the spongin network intact.

### Genotyping

I extracted DNA from some samples with the Qiagen Blood & Tissue kit, and used the Qiagen Powersoil kit on others; downstream results did not differ based on the kit used. At the cox1 locus, a ~1200 bp fragment was amplified with the following primers (LCO1490: 5’-GGT CAA CAA ATC ATA AAG AYA TYG G-3’; COX1-R1: 5’-TGT TGR GGG AAA AAR GTT AAA TT-3’); these amplify the “Folmer” barcoding region and the “co1-ext” region used by some sponge barcoding projects (Rot *et al*. 2006).

Three primer sets were used to amplify portions of the 28S rDNA nuclear locus. Samples were sequenced over the ~800 bp D3-D5 region using primers Por28S-830F (5’- CAT CCG ACC CGT CTT GAA -3’) and Por28S-1520R (5’- GCT AGT TGA TTC GGC AGG TG -3’) (Morrow *et al*. 2012). Most samples were also sequenced over the ~800 bp D1-D2 region using primers Por28S-15F (5’-GCG AGA TCA CCY GCT GAA T-3’) and Por28S-878R (5’-CAC TCC TTG GTC CGT GTT TC-3’) (Morrow *et al*. 2012). A few samples were sequenced using primers C2 (5’-GAA AAG AAC TTT GRA RAG AGA GT-3’) and D2 (5’-TCC GTG TTT CAA GAC GGG-3’) (Chombard *et al*. 1998). The C2-D2 region is a ~50% subset of the D1-D2 region, including the most rapidly evolving region recommended by the sponge barcoding project.

PCR was performed using a Biorad thermocycler (T100); the following conditions were used for the cox1 locus: 95°C for 3 min, followed by 35 cycles of 94°C for 30 sec, 52°C for 30 sec, 72°C for 90 seconds, followed by 72°C for 5 minutes. The 28S C2-D2 region was amplified with the same conditions, except a 57°C annealing temperature and 60 second extension time; the 28S D1-D2 and D3-D5 regions used a 53°C annealing temperature and 60 second extension time. PCR was performed in 50 μl reactions using the following recipe: 24 μl nuclease-free water, 10 μl 5x PCR buffer (Gotaq flexi, Promega), 8 μl MgCl, 1 μl 10mM dNTPs (Promega), 2.5 μl of each primer at 10 μM, 0.75 bovine serum albumin (10 mg/ml, final conc 0.15 mg/ml), 0.25 μl Taq (Gotaq flexi, Promega), 1 μl template. ExoSAP-IT (Applied Biosystems) was used to clean PCRs, which were then sequenced by Functional Biosciences using Big Dye V3.1 on ABI 3730xl instruments. PCR products were sequenced in both directions, and a consensus sequence was constructed using Codon Code v.9 (CodonCode Corporation). Blastn was used to verify that the resulting traces were of sponge origin. All sequences have been deposited in Genbank; accession numbers are listed in table 1.

### Genetic analysis

I used the NCBI taxonomy browser to compile data from all samples identified as belonging to the order Scopalinida. I also used blastn to search for additional sequences that appeared to fall within this order, but found none. At the 28S locus, data from Genbank was only included in the phylogeny if the highly variable C2-D2 region was included. Sequences at cox1 were included if they contained the Folmer barcoding region. Together, included data are from 18 different publications (Blanquer & Uriz 2008; Erpenbeck *et al*. 2002, 2006, 2007a, b, 2012, 2016; Kandler *et al*. 2019; Lavrov *et al*. 2005; Lavrov & Lang 2005; Montalvo & Hill 2011; Morrow *et al*. 2012, 2013; Nichols 2005; Pett & Lavrov 2015; Riesgo *et al*. 2013; Rot *et al*. 2006; Thacker *et al*. 2013). Sequence alignments were produced in Codon Code v.9 (CodonCode Corporation). Phylogenies were estimated with maximum likelihood using IQ-Tree (Nguyen *et al*. 2015; Trifinopoulos *et al*. 2016). Phylogenies are unrooted, and root placement in figures is based on a published phylogeny that had many more characters and representatives from more sponge orders, but limited taxon sampling within the Scopalinida (Morrow *et al*. 2013). I used the Ultrafast bootstrap (Hoang *et al*. 2018) to measure node confidence. Phylogenies were produced from the IQ-Tree files using the Interactive Tree of Life webserver (Letunic & Bork 2019). Figures were made ready for publication using R (r-project.org) and/or Gimp (gimp.org).

## Results

### Genetic Results

Figure 1 shows the phylogentic tree of newly collected samples, previously sequenced Scopalinida, and outgroups at the cox1 mitochondrial locus. The four new species from California form a clade with five previously sequenced *Scopalina*, with *Svenzea zeai* as the closest outgroup. No other closely related sequences could be found in Genbank, so these data support inclusion of the new species in the genus *Scopalina*. Figure 2 shows the phylogentic tree at the 28S nuclear locus (the large ribosomal subunit), which is entirely congruent with results at cox1.

**Figure 1.**
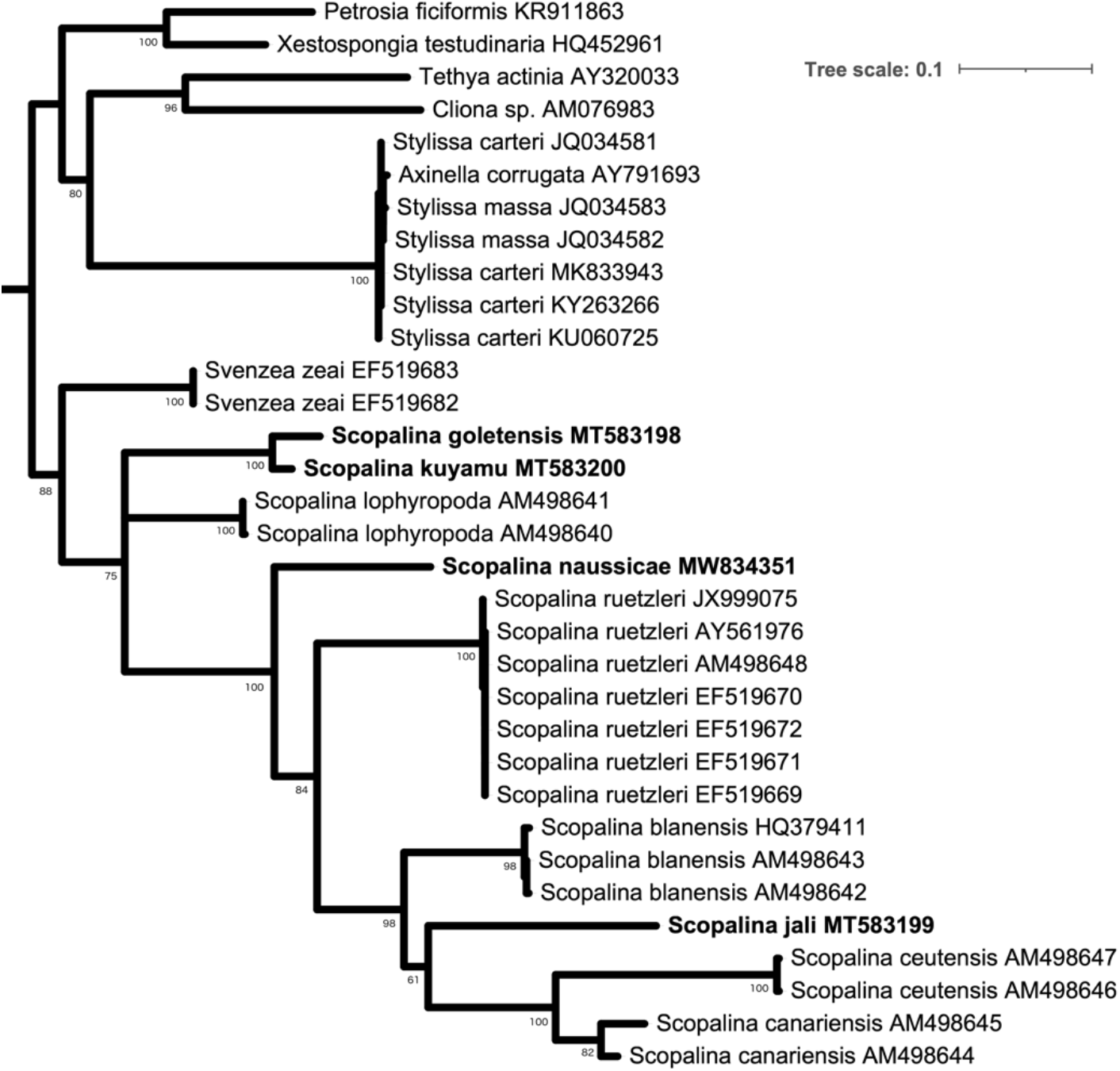
Gene tree at the cox1 mitochondrial locus. Bootstrap values are shown for nodes with > 50% support; nodes with < 50% support are collapsed. Genbank accession numbers are shown; those beginning with M, shown in bold, are new. Scale bar indicates substitutions per site. Root placement based on Redmond et al. (2013).

**Figure 2.**
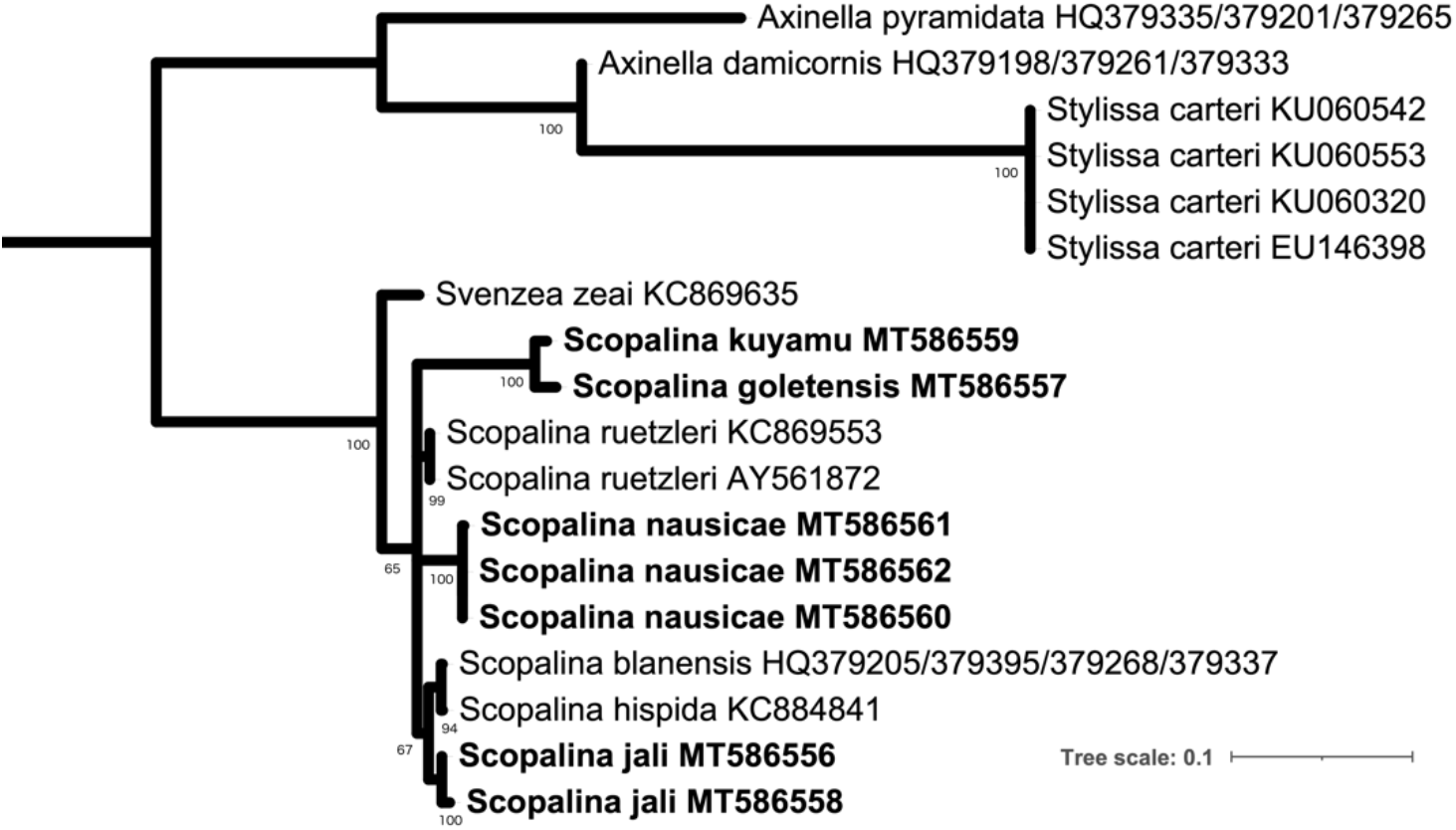
Gene tree at the large ribosomal subunit (28S). Bootstrap values are shown for nodes with > 50% support; nodes with < 50% support are collapsed. Genbank accession numbers are shown; those beginning with MT, in bold, are new. Scale bar indicates substitutions per site. Root placement based on Redmond et al. (2013).

Both gene trees place two of the new species, *S. kuyamu* and *S. goletensis*, as the closest relatives to each other. This raises the question of whether these two species may in fact be a single species. However, these vouchers are different at 4.4% of sites at the cox1 locus, and 3.2% of sites at the 28S locus. The magnitude of this difference is similar to species-level divergence among other Porifera. A review of genetic distances at cox1 found an average 4.9% sequence divergence between any individual sponge sequence and the most closely related sequence available from another species (N=57, (Huang *et al*. 2008)). A more recent analysis of 39 species in the poriferan order Suberitida found 3.7% divergence at this same locus (Turner 2020a).

Together with the morphological differences detailed below, these data support species status for both *S. kuyamu* and *S. goletensis*.

### Systematics

> — Order Scopalinida Morrow & Cárdenas, 2015

#### Definition

Encrusting, massive or erect flabellate growth forms; smooth or conulose surface supported by prominent spongin fibres cored with spicules; megascleres styles and/or oxeas, often with telescoped ends; no ectosomal skeleton; tissue contains an unusual cell type filled with refractile granules. (Modified from Morrow and Cárdenas (2015) to include oxeas among megascleres.)

#### Remarks

As the focus of this paper is alpha taxonomy, rather than a revision of higher taxonomy, I have retained the definition of Morrow and Cárdenas (2015), adding oxeas among the megascleres as the only modification. However, it should be noted that a more thorough revision is needed, and would likely result in further changes. The description of *Svenzea zeai*, for example, does not include prominent spongin fibers.

> — Family Scopalinidae Morrow *et al*., 2012

Definition same as order.

> — *Scopalina* Schmidt 1862

#### Definition

Thinly or thickly encrusting; soft and compressible; few or no ectosomal spicules; spongin abundant, with extensions of spongin manifest as mounds or fibers arising from basal spongin plate; these fibers may branch and merge; choanosomal skeleton of spicules or spicule bundles with proximal ends or entire spicule enclosed in spongin; choanosome may have a grainy appearance. Larvae are elongated, conical; anterior region wider than the posterior zone; completely covered by short cilia. (Modified from Blanquer and Uriz 2008).

#### Diagnosis

*Scopalina* have abundant spongin, while *Svenzea* are described as having limited spongin, primarily at the nodes of a reticulated spicule network. *Svenzea* tend to have shorter spicules, (200-300 μm), whereas in *Scopalina* they mostly range from 400 to 2000 μm (though *S. canariensis* averages only 199 μm). The skeletal architecture of *Svenzea* has been noted as more like that of the haplosclerida than *Scopalina*. *Svenzea* are massive or thickly encrusting, while *Scopalina* are thinly to thickly encrusting.

*Stylissa* are erect, flabellate, or lobate, rather than possessing encrusting morphologies seen in *Scopalina*. *Stylissa* are noted as having a skeletal architecture like that of the Halichondridae, with many spicules in confusion.

> — *Scopalina nausicae* **sp. nov.**

#### Material examined

Holotype: (CASIZ 235474) Point Loma, San Diego, California, USA (32.69438, −117.26860), 15 m depth, 2/7/20. Paratypes: (CASIZ 235471) Coal Oil Point, Santa

Barbara, California, USA (34.40450, −119.87890), 11 m depth, 8/30/19; (CASIZ 235472) Isla Vista Reef, Santa Barbara, California, USA (34.40278, −119.85755), 12 m depth, 8/1/19; (CASIZ 235473) Arroyo Quemado Reef, Santa Barbara, California, USA (34.46775, −120.11905), 11 m depth, 1/7/20.

#### Etymology

Named for the fictional character Nausicaä from the film *Nausicaä and the Valley of the Wind*.

#### Morphology

Encrusting, 2 - 4 mm thick, up to 10 cm across (Fig. 3). Soft and compressible. Prominent conules 0.5 - 1.0 mm in height,1.5 - 3.5 mm apart; spicules protrude at conules, making them microscopically hispid. Scattered oscules 1 - 2 mm in diameter. In nature, ectosome appears opaque at conules but often lacy and porous between them; ectosome more opaque in collected samples. Ectosome peach colored, choanosome yellow when alive; all tissues fade to beige when preserved in ethanol.

**Figure 3.**
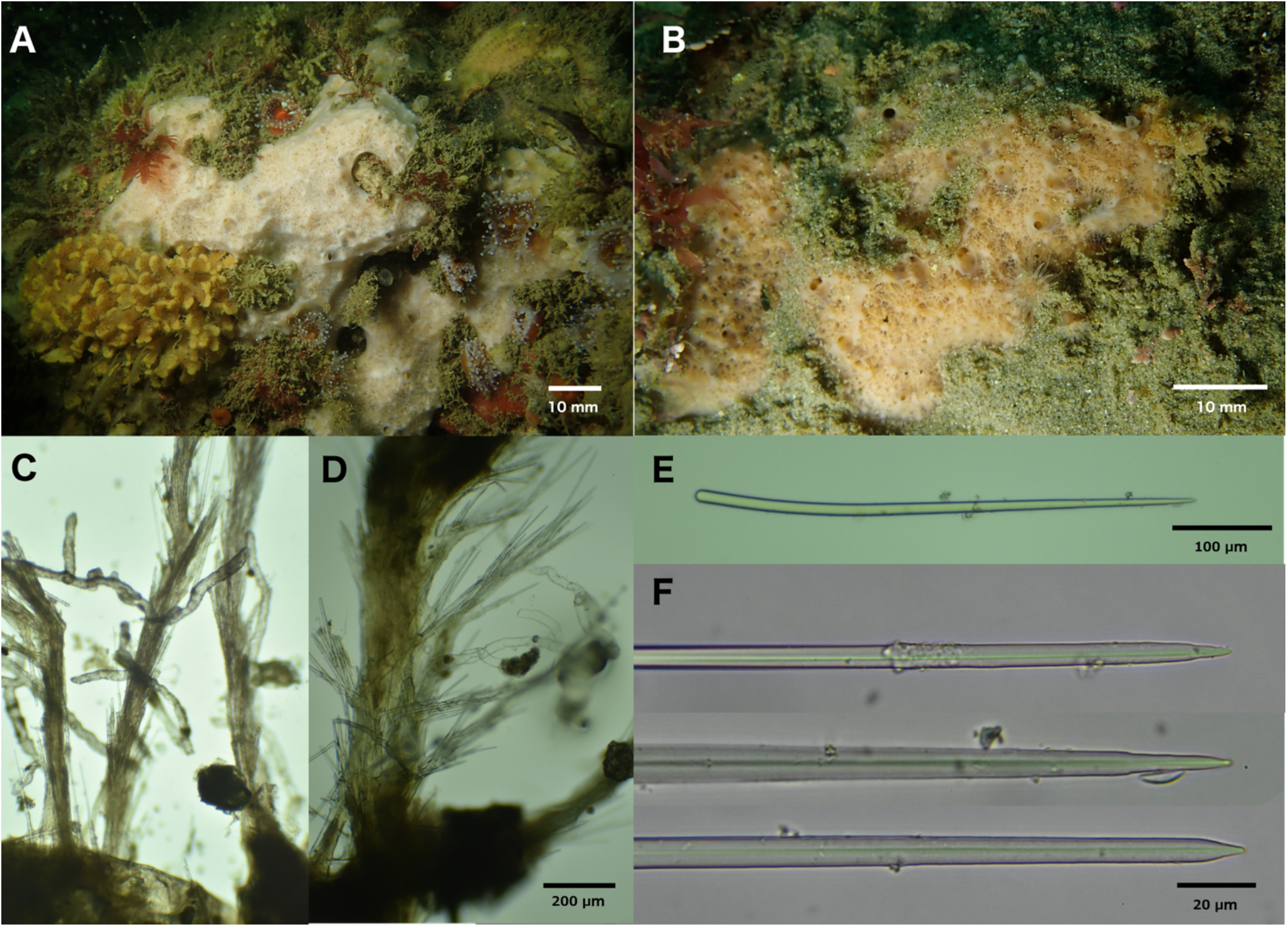
*Scopalina nausicae* **sp. nov**. A: Field photo of paratype (CASIZ 235472), Isla Vista Reef. B: Field photo of paratype (CASIZ 235473), Arroyo Quemado Reef. C, D: Skeletal architecture of holotype (CASIZ 235474), Point Loma; scale applied to both. In C, vermiform and primary spongin trunks are coated in apparent algae. In D, only primary spongin trunk showing algae, but vermiform tracts are sporadically cored with sediment. E, F: spicules from paratype (CASIZ 235472).

#### Skeleton

Vertical trunks of spongin, 100 - 550 μm wide, arise from a basal spongin mat and terminate in surface conules. Secondary branches of spongin 50 - 100 μm wide arise from primary trunks, branching off at an angle of less than 90 degrees and still extending towards surface. Primary and secondary trunks cored with spicules with pointed ends up; spicules entirely enclosed in spongin or with tips projecting; projecting tips fan out to create a bouquet that pierces the ectosome at conules. An additional type of spongin tract is distinct from those described above: 60 - 90 μm wide, these tracts branch from primary trunks at approximately 90-degree angles, then meander through the choanosome in a vermiform fashion, sometimes branching; these vermiform tracts do not contain spicules. Basal spongin, spicule-containing spongin trunks, and vermiform tracts are sporadically cored with sediment. Spicule-containing and vermiform spongin tracts are often filled and/or coated with what appear to be algal cells; these are red in preserved tissue.

#### Spicules

Styles only, usually bent towards the head end, thickest at the head and tapered to a point. Some show “telescoping” (width decreasing in a step-wise fashion) at the pointed end. Average spicule length for each voucher: 454, 483, 505, 532 μm (N=31-40 per sample); total range in spicule length across vouchers 375-623 μm (N=135). Average spicule width at head, for each voucher: 9, 9, 11, 11 μm (N=31-40 per sample); total range in spicule width at head 5-17 μm (N=135).

#### Distribution and habitat

This species is common on the shallow (5-16 m) rocky reef at Coal Oil Point, Santa Barbara, California. Often found on vertical rock walls or boulders, it can also occur on flatter areas, and has been found partially buried by sand. It was not found at most other locations investigated, but was located in similar habitat at the Arroyo Quemado Reef (near Point Conception) and in the kelp forests in extreme Southern California, off Point Loma and La Jolla, San Diego. It is therefore likely that the specie’s range encompasses at least the Southern Californian and Ensenadan biogeographical provinces, bounded by Point Conception in the North and Punta Eugenia in the South (Blanchette *et al*. 2008; Valentine 1966).

#### Remarks

Skeletal architecture, spiculation, and genotype all conspire to place this species within the *Scopalina*. I was unable to detect the “graininess” said to characterize other Scopalinidae. However, this was hard to assess due to the abundant sediment within the sponge: dark grains were apparent, but appeared to be sediment rather than refractile cells.

Spicule dimensions, skeletal morphology, and genotype all serve to differentiate *S. nausicae* **sp. nov**. from the three other species newly described here. Fourteen other species are currently placed in the genus *Scopalina*, according to the World Porifera Database (van Soest *et al*. 2019). None of these are known from the Eastern Pacific, making them unlikely conspecifics with any of the species described here. The gross morphology of *S. nausicae* **sp. nov.** in the field is quite similar to published images of *S. ruetzleri* (Wiedenmayer, 1977) (West Atlantic) and *S. erubescens* (Goodwin *et al*., 2011) (Faulkland Islands). Spicule length and sponge color also match *S. erubescens* better than other *Scopalina*, making this species the most likely conspecific. In addition to geographic separation, however, *S. erubescens* is larger, more thickly encrusting, and has thicker spicules and spicule bundles. The description of *S. erubescens* also lacks any mention of the vermiform spongin tracts that pervade *S. nausicae* **sp. nov**. (Goodwin *et al*. 2011)*. Scopalina ruetzleri* can be excluded as a conspecific based on genetic data at both cox1 and 28S as well as color and habitat (Rützler *et al*. 2003). This species is described as ranging throughout the Caribbean, but was also recently reported from the tropical Eastern Pacific (Carballo *et al*. 2019). This latter report is not accompanied by morphological or genetic information, so comparisons between tropical Pacific *Scopalina* and *S. nausicae* **sp. nov**. await future investigation.

Within its range, it is likely that this sponge can be identified from field photos, as I have seen no other sponge with a similar morphology to date.

> — *Scopalina kuyamu* **sp. nov**.

#### Material examined

Holotype: (CASIZ 235469) Naples Reef, Santa Barbara, California, USA (34.42212, −119.95154), 12 m depth, 7/31/19.

#### Etymology

Named for the village of Kuyamu, a community of Barbareño Chumash that once stood onshore at the site where the sponge was discovered.

#### Morphology

Encrusting, 1-2 mm thick, 6 cm across (Fig. 4). Soft and compressible. Surface hispid due to a profusion of protruding styles. Distinct ectosome not apparent. Peach colored in nature, except for translucent-white varicose channels running along surface. Few scattered oscules, each ~300 μm diameter; smaller pores (approximately 80 μm diameter) abundant and uniformly distributed. Beige when preserved in ethanol.

**Figure 4.**
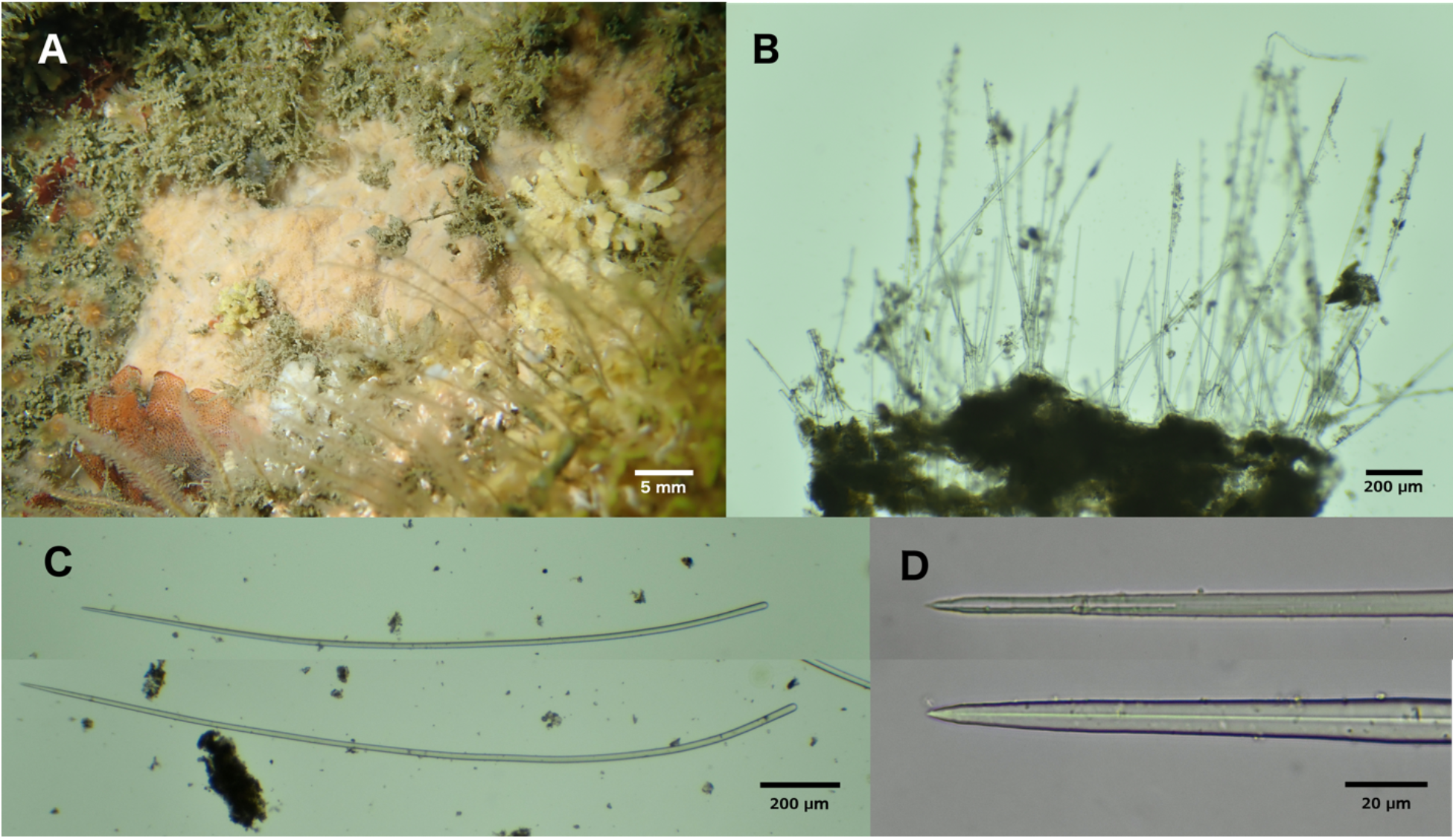
*Scopalina kuyamu* **sp. nov**. All images from holotype. A: field photo. B: Skeletal architecture, showing basal plate cored with copious sediment. C, D spicules.

#### Skeleton

Basal mat of spongin cored with sediment. Extensions of spongin arise from this mat: most are low mounds, some only 25-50 μm high, but some are fingers 100-300 μm high and cored with sediment. Heads of spicules are embedded in these mounds and fingers, either singly or in bundles of up to 12. Spicules extent vertically and pierce the surface of the sponge.

#### Spicules

Styles only, usually bent towards the head end, thickest at the head and tapered to a point. Some spicule tips are “telescoping” (width decreasing in a step-wise fashion) at the pointed end. Spicules averaged 1557 μm in length (N=35, range 879-1948 μm); 16 μm in width (N=35, range 11-21).

#### Distribution and habitat

Only a single individual has been found, on a vertical wall at 12 m depth, at Naples Reef, in Santa Barbara, California. Habitat was rocky reef with abundant bryozoan, sponge, and anthozoan cover, adjacent to year-round kelp forest. Three additional dives at the same location failed to locate other individuals; similar, nearby habitat to the East and West also had considerable search effort, so this species appears to be rare in this area.

#### Remarks

This sponge is quite genetically and morphologically distinct from *S. nausicae* and *S. jali*. The spicular architecture is fairly similar to *S. goletensis*, though the spicule density is lower. As a result, *S. goletensis* is removable from the substrate as a fairly firm sheet, while *S. kuyamu* peels away in rubbery strips that curl up upon themselves. Also, the spicules average over twice as long in *S. kuyamu*, with non-overlapping size ranges among the spicules measured. These morphological differences seem unlikely to be due to environmental influences, as the two species were collected at the same depth, at very similar reefs, less than 5 km apart. Together with the considerable genetic divergence, these differences support species status for both species.

Among *Scopalina* from other regions, the only species with spicules as large as *S. kuyamu* are *S. lophyropoda* (Schmidt, 1862) (Blanquer & Uriz 2008) (Mediterranean) and *S. bunkeri* (Goodwin *et al*., 2011) (Falkland Islands). In addition to great geographic distance, *S. lophyropoda* can be excluded based on genetic data (Fig. 1); *S. bunkeri* has a different spicular architecture, gross morphology, and color (Goodwin *et al*. 2011).

It does not seem likely that this species can be identified from field photos alone, though it is difficult to say if there are reliable field marks until more individuals are found.

> — *Scopalina goletensis* **sp. nov**.

#### Material examined

Holotype: (CASIZ 235470) Elwood Reef, Santa Barbara, California, USA (34.41775, −119.90150), 12 m depth, 10/23/19.

#### Etymology

Named for the town of Goleta that is onshore from the location where the sponge was discovered.

#### Morphology

Encrusting, 1.0 - 1.2 mm thick, approximately 2.5 cm across (figure 5). Firm and incompressible. Surface hispid due to dense profusion of protruding styles. Distinct ectosome not apparent. Beige / cream colored in nature; retained the same color when preserved in ethanol.

Surface traced by varicose, translucent channels; pores (approximately 200-300 μm diameter) abundant and uniformly distributed.

**Figure 5.**
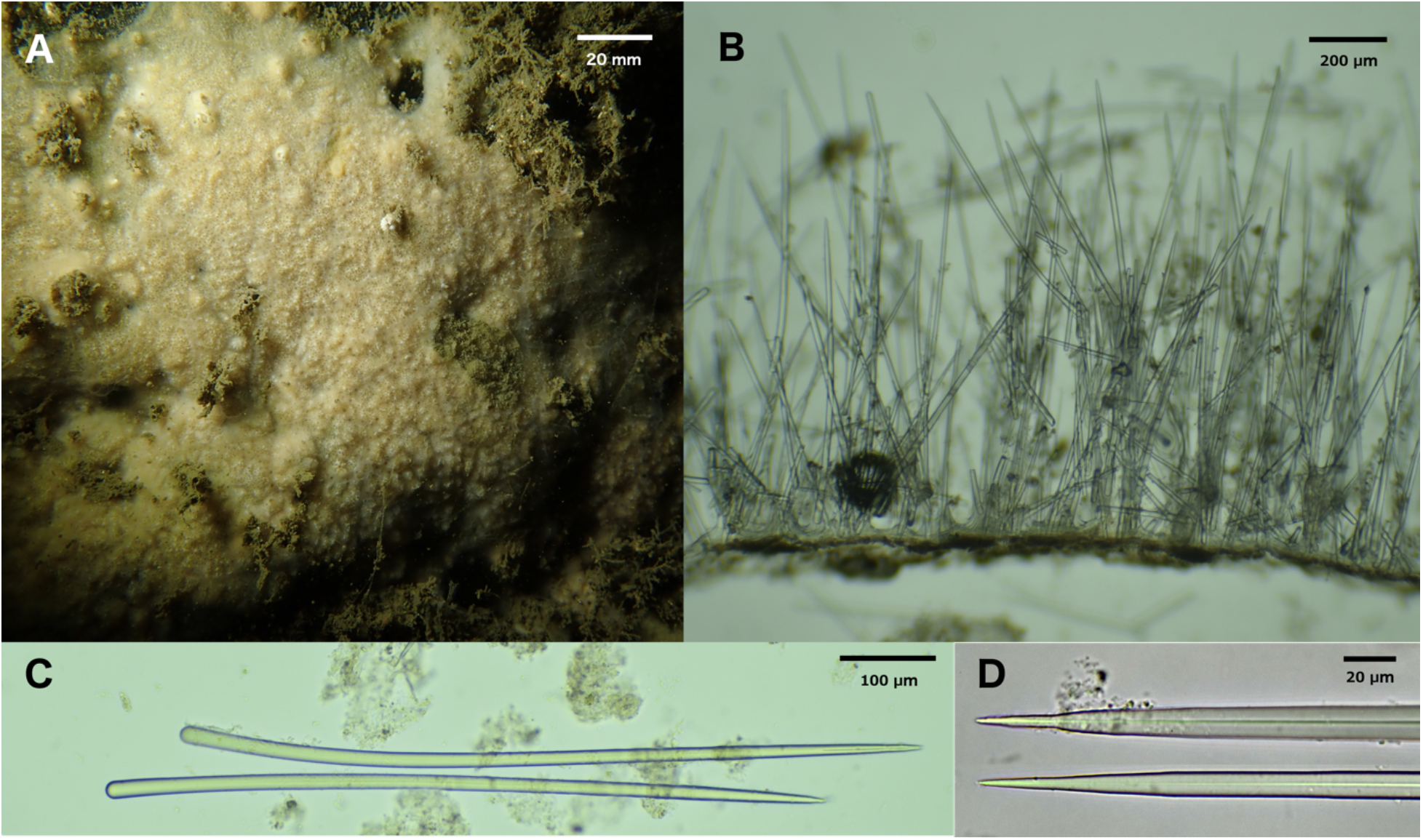
*Scopalina goletensis* **sp. nov**. All images from holotype. A: field photo. B: Skeletal architecture, showing basal plate cored with sediment. C, D spicules.

#### Skeleton

Basal mat of spongin cored with sediment. Vertical extensions of spongin 10-600 μm high arise from this mat: none of these were cored with sediment, but loose sediment was abundant throughout the sponge. Heads of some spicules are embedded singly, directly in the basal mat of spongin, but most are embedded as tiered bundles in the vertical extensions of spongin.

#### Spicules

Styles only, usually slightly bent towards the head end, thickest at the head and tapered to a point. Some spicule tips are “telescoping” (width decreasing in a step-wise fashion) at the pointed end. Spicules averaged 687 μm in length (N=37, range 388-801 μm); 15 μm in width (N=37, range 6-21).

#### Distribution and habitat

Only a single individual has been found, on a vertical ledge at 12 m depth, at Elwood Reef, in Santa Barbara, California. Habitat was rocky reef with abundant bryozoan, sponge, and anthozoan cover, under a year-round kelp canopy. Considerable search effort at Elwood Reef and nearby locations failed to locate additional individuals, so this species is likely to be rare in this area.

#### Remarks

This species is most similar to *S. kuyamu*, but is morphologically and genetically distinct, as detailed in the *S. kuyamu* remarks. The spicule dimensions are similar to several species from other regions (*S. azurea* (Bibiloni, 1993), *S. blanensis* (Blanquer & Uriz, 2008), *S. hispida* (Hechtel, 1965)), though none of these others is known to have spicules as thick. All but *S. azurea* can also be excluded based on the available genetic data (Figs. 1, 2). Conspecificity with *S. azurea* is unlikely based on geographic isolation, color, and spicular architecture (Bibiloni 1993).

It does not seem likely that this species can be identified from field photos alone, though it is difficult to say if there are reliable field marks until more individuals are found.

> — *Scopalina jali* **sp. nov**.

#### Material examined

Holotype: (CASIZ 235466) Big Rock, Santa Cruz Island, California, USA (34.05220, −119.57360), 12m depth, 1/19/20. Paratypes: (CASIZ 235467) Naples Reef, Santa Barbara, California, USA (34.42212, −119.95154), 11 m depth, 9/26/19. (CASIZ 235468) Naples Reef, Santa Barbara, California, USA (34.42212, −119.95154), 15 m depth, 12/10/19.

#### Etymology

The ectosome of live specimens *in situ* is reminiscent of a jali: a latticed screen common in Indo-Islamic architecture.

#### Morphology

Thickly encrusting, 1.0 - 1.5 cm thick, up to 35 cm across (Fig. 6). Soft, spongey, and very compressible. Ectosome transparent, without spicules; a lattice-like mesh of spongin fibers visible in life; conules present, but very small (100-300 μm in width and height); ectosome more opaque after preservation in ethanol but remains partially transparent and lacy. Color in freshly collected specimens is terra-cotta (reddish-brown); red and orange tones are more muted in field photos, with color appearing to vary from tan to terra-cotta; samples fade to beige when preserved in ethanol. Oscules 10 - 20 mm in diameter; occur singly; sparse in some samples and denser in others; partially closed by ectosomal membrane in collected samples.

**Figure 6.**
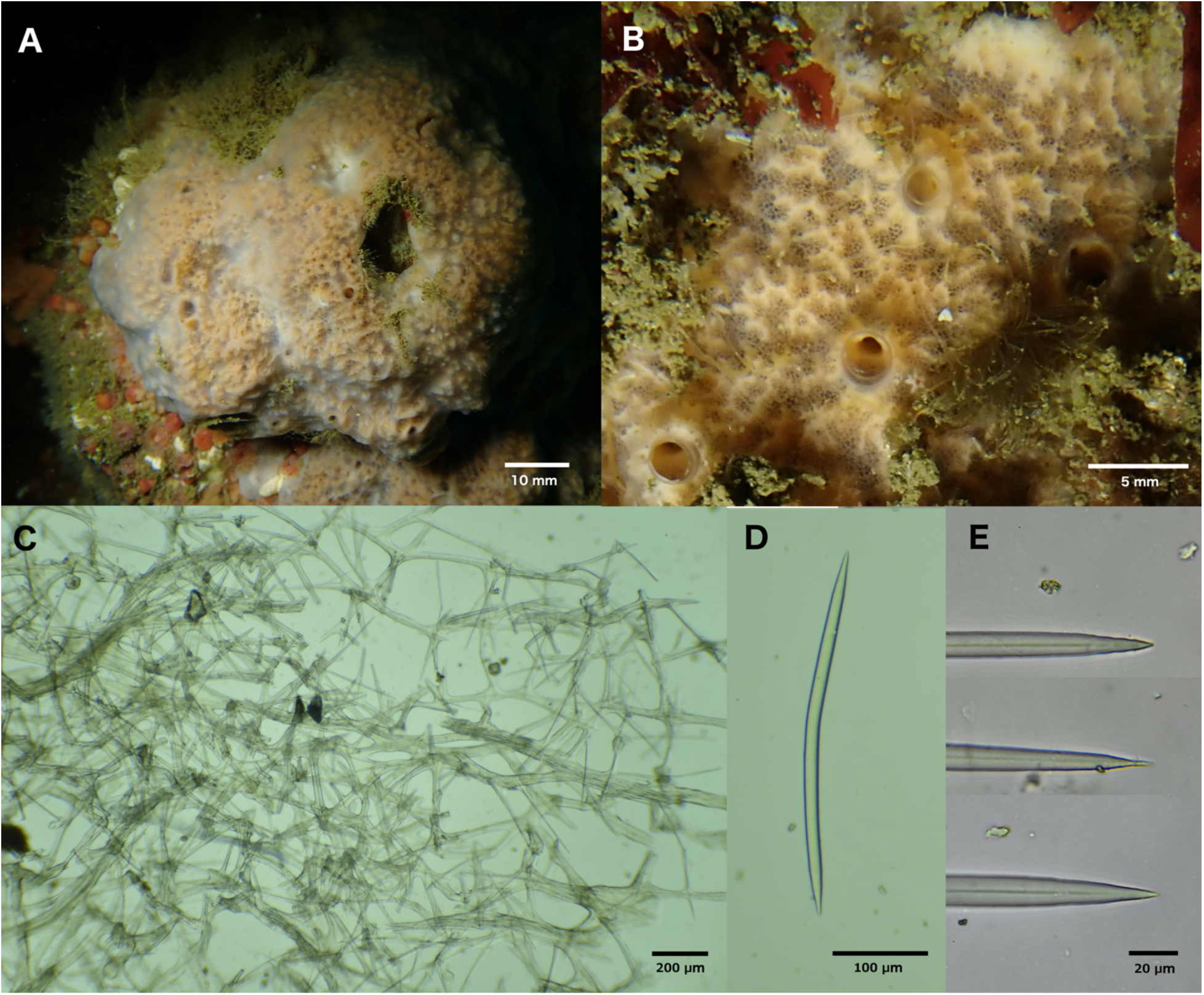
*Scopalina jali* **sp. nov.**. A: Field photo of holotype (CASIZ 235466), Santa Cruz Island. B: Field photo of paratype (CASIZ 235467), Naples Reef. C: Skeletal architecture of holotype. In C, vermiform and primary spongin trunks coated in apparent algae. D, E: spicules from paratype (CASIZ 235467).

#### Skelseton

Abundant spongin fibers cored with spicules form a chaotic mesh lattice within choanosome. Larger spongin tracts, 45 - 65 μm wide, are cored with bundles approximately 5 spicules wide; smaller tracts, 8 - 20 μm wide, are cored with single spicules. No spicules detected outside of spongin tracts. Considerable silt apparent in tissue sections, but none seen coring spongin tracts.

#### Spicules

Oxeas only, gently curved; some spicule tips show “telescoping” (width decreasing in a step-wise fashion). Average spicule length for each voucher: 354, 358, 366 μm (N=30-37 per sample); total range in spicule length across vouchers 219-436 μm (N=100). Average spicule width at widest point, for each voucher: 8, 9, 11 μm (N=30-37 per sample); total range in spicule width at head 2-18 μm (N=100).

#### Distribution and habitat

In the winter of 2019-2020, this sponge was abundant on the shallow (5-17 m) rocky reefs off of Naples Point and the Elwood Bluffs, Santa Barbara, California. The species was not seen in 4 dives at these same locations in the Spring and Summer of 2019, suggesting that the population may vary seasonally or in a boom-and-bust fashion on longer timescales. Consistent with this latter possibility, many large individuals of this species were seen at the Big Rock dive site at Santa Cruz Island in January of 2020, while no individuals were seen in three dives at the same location in November of 2018. The only other probable sighting I am aware of is a photo uploaded to the site iNaturalist (inaturalist.org/observations/41000570). This photo is very likely to be *S. jali*, as no other sponge with this morphology is known in Southern California. The photo is annotated as from Heisler Park, Laguna Beach, from 3/4/2007.

#### Remarks

Genetic data at two loci confirm that this species is within the *Scopalina*. Abundant spongin fibers cored with simple spicules, telescoping spicule tips, and lack of ectosomal skeleton are all consistent with this placement. The presence of oxeas, rather than styles, required modification of recent definitions of order, family, and genus -- though one species currently placed in *Scopalina* in the World Porifera database also contains only oxeas (*S. agoga* (de Laubenfels, 1954)) and another contains both styles and oxeas (*S. australiensis* (Pulitzer-Finali, 1982)). *Scopalina jali* is differentiated from *S. agoga* by spicule size and the presence of many tangential spicules in the ectosome of *S. agoga*; this previously described species is also known only from Palau (de Laubenfels 1954). The skeletal architecture of *S. jali* differs markedly from the other California species described herein due to its highly reticulated nature, but this is similar to the published description of the Atlantic species *S. ceutensis* (Blanquer & Uriz, 2008).

It is likely that this sponge can be identified from field photos alone within Southern California, as I have seen no other sponge with a similar morphology to date.

## Conclusions

It is remarkable that, among the relatively well-studied kelp forests of the Santa Barbara Channel, I was able to locate 4 new species from an order not previously known to occur in the region. These species were not only undescribed, but apparently unsampled: no previous California survey includes samples matching their description (Lee *et al*. 2007). These results illustrate how much remains to be learned about the sponges of California, and show that SCUBA-based collection efforts can help bridge this gap. Moreover, collection by hand allows for underwater photography of live samples before collection. By comparing photographs of the species described here with photos of all other sponges I have collected, I believe that the two more common species (*S. jali* and *S. nausicae*) are easily recognizable within their range. This assertion is supported by the fact that, after collection of the first samples of each, subsequent samples were correctly identified as conspecifics based on field photos before being confirmed as such using spicules and DNA. This will simplify future efforts to understand the ecology of these species, perhaps by using diver surveys or photo transects.

In contrast, the other two new species (*S. kuyamu* and *S. goletensis*) are thinly encrusting and will be harder to identify based on gross morphology. Each was found only once; as they were both found within the range and habitat I have most thoroughly sampled, it is likely they are uncommon in this area (but could be common in deeper waters). Though describing new species from a single sample is not ideal, I feel confident these samples are not conspecific with any currently named species due to their considerable genetic divergence from other sequenced species, substantial morphological differences from un-genotyped species, and the fact that no previously named species are known from the region.

Much remains to be learned about the Scopalinida. Twenty species were previously known to reside in the order: 14 *Scopalina* species, 5 *Svenzea* species, and *Stylissa flabelliformis* (Morrow & Cárdenas 2015). These sponges are known from the Mediterranean and Canary Islands (5 species), Caribbean (6 species), the tropical South-West Pacific (6 species), Madagascar (1 species), and the Falkland Islands (2 species). The addition of four species from Southern California expands this range considerably, and makes the group more accessible to researchers in this geographic region. It is my hope that this will lead to an improved understanding of the ecology and evolution of these interesting and understudied animals.

## Acknowledgements

I am grateful for the help and support of many people in UCSB’s Marine Science Institute and Diving & Boating Program, especially Robert Miller, Clint Nelson, Christoph Pierre, Frankie Puerzer, Christian Orsini, and H. Mark Page. Steve Lonhart (NOAA) and Shannon Myers (UCSC) were instrumental in facilitating collections in Central California, and the Natural History Museum of Los Angeles’ DISCO program facilitated collections in Los Angeles County. I’d also like to thank the editor and two anonymous reviewers for their help in improving the paper.

